# Small phytoplankton community composition cycles annually with a coastal bloom

**DOI:** 10.1101/2025.09.15.675855

**Authors:** Bethany L. F. Stevens, Rebecca J. Gast, Emily E. Peacock, Yogesh A. Girdhar, Michael G. Neubert, Heidi M. Sosik

## Abstract

Small photosynthetic eukaryotes are a productive and dynamic component of marine planktonic communities. Here, we investigate how seasonal changes in abundance of these primary producers relate to changes in their community composition at a coastal site on the Northeast U.S. Shelf. We present a 9-yr time series of 18S rRNA sequencing data and identify gradual transitions within the pico- and nanoplankton community that occur repeatedly over the annual cycle. We compare these compositional changes to concurrent high-resolution in situ flow cytometry measurements of eukaryotic phytoplank-ton abundance and division rate. We find that the Chlorophyta contribute a large proportion of the sequences in our samples and drive much of the seasonal variability within the small phytoplankton community. Across the time series, *Bathycoccus, Micromonas*, and *Picochlorum* are the dominant genera, with the first being present year round, while *Micromonas bravo* and *Picochlorum* are representative of the summer community. We also find a strong winter *Phaeocystis* signal which might be leading to flow cytometry measurements of relatively large cells in the early spring. Our results provide fundamental knowledge of the taxonomic composition of the phytoplankton community on the Northeast U.S. Shelf, improving our understanding of the region’s diversity and compositional variability over time.

## 1 Introduction

Marine phytoplankton make up less than 1% of Earth’s photosynthetic biomass, but are responsible for almost half of all primary production (Field et al., 1998). Among phytoplankton, the pico- and nano-sized eukaryotes are particularly extreme in their production relative to their size and abundance. Too small to be identified through microscopy and less abundant than cyanobacteria, the small eukaryotes are notable for their rapid growth rates and high turnover in the planktonic food web (Li, 1994; Worden et al., 2004; Fowler et al., 2020). For example, Worden et al. (2004) estimated that a biomass equal to 250% of the standing stock was consumed and replaced through growth on a daily basis.

The small eukaryotes are among the plankton that dominate marine waters under warm, oligotrophic conditions. In temperate regions, spring blooms can occur when conditions shift in favor of these smaller cells (O’Reilly and Zetlin, 1998; Tamigneaux et al., 1999). Individual cells can be measured through flow cytometry on the basis of their autofluorescence and light-scattering signatures, but the phytoeukaryotes smaller than 10*µ*m cannot be be taxonomically classified through flow cytometry alone. High-throughput sequencing and metabarcoding have shown the small phytoeukaryotes to be incredibly taxonomically diverse (Moon-van Der Staay et al., 2001; D’iez et al., 2001; Zeidner et al., 2003). It is rarely possible, therefore, to attribute changes in their abundance to a particular taxa, and many questions remain as to how the ecology of the group depends upon its taxonomic composition.

On the Northeast U.S. Shelf, the small phytoplankton community undergoes a dramatic annual spring bloom (O’Reilly and Zetlin, 1998; Hunter-Cevera et al., 2016). Since 2003, phytoplankton at the Martha’s Vineyard Coastal Observatory (MVCO) have been detected and measured with automated flow cytometry at a twenty minute resolution (Olson et al., 2003; Sosik and Olson, 2022). These data have been used to monitor cell abundance as well as to estimate daily cell division rate (Hunter-Cevera et al., 2014; Hunter-Cevera et al., 2019; Fowler et al., 2020). Moreover, MVCO is open to advection, and the ecological information gained from the site has been shown to be relevant for small phytoplankton across much of the Northeast U.S. Shelf (Stevens et al., 2023, 2024). While aggregate properties of the small phytoeukaryotes at MVCO have been closely monitored for decades, no corresponding information about taxonomy has been reported. We therefore set out to i) determine which taxa contribute to the small phytoplankton assemblage at MVCO, ii) assess the temporal variability of this community’s taxonomic composition and relate it to changes in cell abundance and division rate, and iii) identify groups of taxa that co-occur regularly and may warrant future study of competitive or other interaction.

With the above objectives in mind, we collected a 9-y time series of monthly genetic samples from surface waters at MVCO. While amplicon sequencing is a powerful method for detecting taxa, metabarcoding data are only semiquantitative (Gloor et al., 2017). We therefore leverage the flow cytometry time series to complement our analysis with quantitative measures of cell abundance. We also implement a suite of analytical techniques that handle compositional genetic data appropriately. These methods include PCA for characterizing variability, SparCC (Sparce Correlations for Compositional Data) for identifying correlations between pairs of taxa (Friedman and Alm, 2012), and spatiotemporal topic modeling for describing our samples as mixes of co-occurring communities (Girdhar et al., 2014). This last tool, though rarely used for genetic data, is well suited for their analysis, and our work demonstrates how it could be adopted for broader use in microbial ecology. Through these combined methods, we find seasonal changes in the composition of the small phytoplankton community that correspond closely to patterns in pico- and nano-eukaryote abundance. These observations allow us to begin partitioning the well-described aggregate dynamics of small phytoplankton on the Northeast U.S. Shelf into the responses of individual taxa.

## 2 Methods

### 2.1 Sample collection and processing

The MVCO offshore tower (41^*°*^19.500’ N, 70^*°*^34.0’ W) is the most nearshore station of the Northeast U.S. Shelf Long-Term Ecological Research site. The tower is *∼*2 km south of the island of Martha’s Vineyard where the water column has a depth of 15 m and is typically well-mixed by winds and tides. The predominant regional currents flow from the east along the continental shelf (Limeburner and Beardsley, 1982). Genetic samples for metabarcoding were collected from the upper 4 m of seawater at MVCO by either bucket or Niskin bottle roughly monthly between February 2013 and December 2021. Samples were filtered and prepared for sequencing as described in Catlett et al. (2023): for each sample, between 700 and 1000 ml of whole seawater were filtered onto a 0.2 *µ*m filter. Cells from one half of each filter were lysed and their nucleic acids extracted with the ZYMO Quick-DNA Fungal/Bacterial Microprep Kit. The V4 region of the 18S ribosomal RNA gene was targeted for amplification with the 574*f and 1132r primers described in Hugerth et al. (2014). These primers were developed specifically to resolve eukaryote diversity, but their length creates a challenge for merging forward and reverse reads. Only forward reads were included in our analysis, as in previous work with the same primers (Millette et al., 2021). Samples were sequenced on an Illumina MiSeq (v3 600; 2 x 300 bp) at the URI Genomics and Sequencing Center (sample dates Feb 2013-Aug 2017 and Aug 2019-Oct 2020) and the Georgia Genomics and Bioinformatics Core (sample dates Aug 2017-Jul 2019 and Nov 2020-Dec 2021).

Amplicon sequence variants (ASVs) were inferred from demultiplexed sequence data using the DADA2 pipeline (Callahan et al., 2016). First, forward reads were trimmed to 190-210 nucleotides based on declines in their quality profiles at higher values. Reads were further truncated if quality scores fell below 10 or if the maximum expected error was greater than 2. These values were chosen to be more conservative than the DADA2 default parameters. As mentioned above, we did not merge forward and reverse reads. Chimeras were detected and removed using the removeBimeraDenovo function’s default “consensus” method. The remaining 18,680,520 sequences were composed of 18,752 unique ASVs for our analysis.

We analyzed the relationships between our metabarcoding data and environmental measurements made on the same days that the genetic samples were collected from MVCO. These include variables measured continuously at the observatory—water temperature and salinity recorded by MicroCat conductivity, temperature, pressure sensor, and incident short wave radiation recorded by an Eppley pyranometer—as well as dissolved nutrients in seawater samples collected from *<*10 m depth (Sosik et al., 2021). Nutrient samples were filtered, frozen, and processed at the Woods Hole Oceanographic Institution’s Nutrient Analytical Facility.

We also compared our compositional genetic data to direct measurements of phytoplankton concentration. Automated flow cytometers, or FlowCytobots, have been deployed in succession at MVCO since 2003. These instruments are fixed 4 m below the surface and sample the surrounding seawater at 20-minute intervals (Olson et al., 2003). Eukaryotic phytoplankton in the pico-(0-2 *µ*m) and small nano-(2-10 *µ*m) size classes have been identified from these measurements based on their light-scattering and fluorescence signals. Here, we supplement the existing time series of in situ measurements with discrete samples collected from *<*10 m depth at MVCO on 50 days between 2015 and 2021. These samples were frozen in liquid nitrogen and later thawed and analyzed in the lab with an Attune NxT Flow Cytometer (ThermoFisher). Our total data set includes genetic samples from 125 days and over 40,000 flow cytometry measurements collected during the same time period (Woods Hole Oceanographic Institution, 2018, 2024; Sosik and Olson, 2022).

### 2.2 Assessing variability in the small phytoplankton community

To identify small phytoplankton taxa, unique ASVs in our dataset were compared to the Protistan Ribososomal Reference (PR2) database v4.14.0 (Guillou et al., 2013). We used the assignTaxonomy function in DADA2 which implements the RDP Naive Bayesian Classifier algorithm (Wang et al., 2007). We then focused our analysis on those taxa likely to be monitored by flow cytometry by considering two subsets of the total genetic data set. The first focal group is restricted to eukaryotic photosynthetic plankton, and further reduced to exclude taxonomic groups strictly outside the pico- and nano-size classes: mixotrophic Radiolarians and photosynthetic members of the class Euglenida were assumed to be above the target cell size. Members of the phylum Dinoflagellata and the class Bacillariophyceae were excluded unless classified at the species level to a species identified as pico- or nano-size by Vaulot et al. (2008) or Mitra et al. (2023). A complete list of the included species is attached as a supplementary file. This first focal group, which we will call “small phytoplankton,” includes 2,528 ASVs within our dataset. Because we do not have size information for every species, this focal group may include some ASVs corresponding to cells which are larger than the cells measured by flow cytometry. We carried out more detailed analysis on ASVs assigned to the phylum Chlorophyta, the majority of which are well within the size range measured by flow cytometry and make up the largest portion of the reads assigned to the small phytoplankton group. Note that this second focal group is a strict subset of the first small phytoplankton group and so does not include chlorophyte macroalgae in the class Ulvophyceae.

From the classified genetic data, we identified temporal patterns in the composition and diversity of the small phytoplankton assemblage. We calculated Shannon diversity at the genus level, excluding ASVs that were assigned to a genus with less than 80% confidence so as not to overinflate proportions of incorrectly assigned taxa. Then, to characterize the variability across samples, we performed principal component analysis (PCA) from species-level taxonomic assignments within each of our two focal groups described above. For these PCAs, we again excluded ASVs assigned with less than 80% confidence and then performed a centered log ratio transformation to the read counts, as recommended by Gloor et al. (2017). We plotted the values of the first two principal components over time to visualize temporal changes in the community.

We also used the results of the PCA to relate variability in the community composition to environmental conditions at MVCO. We measured correlation between principal component values and the available environmental variables.

When correlation was statistically significant (*α* = 0.05), we tested the variable’s predictive ability and compared it to that of day-of-year. In particular, to evaluate whether temperature or day-of-year better explained variation in the small phytoplankton community, we conducted a 10-fold cross-validation experiment on degree-3 polynomial regressions (James et al., 2021). The cubic polynomial was chosen to capture the smooth seasonal cycles in the variables of interest. 90% of the observations were used to train the model for each predictor and the remaining 10% were used to evaluate the model by calculating the mean squared error. We then applied a paired *t*-test to the mean squared errors to determine whether one model consistently outperformed the other.

### 2.3 Identifying co-occurring groups

The final objective of our analysis was to identify groups of small phytoplankton taxa that co-occur regularly at our study site. We used both SparCC inference and spatiotemporal topic modeling to detect pairs and groups of co-occurring taxa, respectively. SparCC is a tool designed for metabarcoding data that infers positive correlations from the assumptions that data are high-dimensional, sparse, and compositional (Friedman and Alm, 2012). Strong positive correlations indicate pairs of individual taxa that co-occur regularly, suggesting a shared niche and the potential for either competitive or facultative relationships. We implemented the SparCC algorithm in the R programming language (R version 4.1.3 (2022-03-10); R Core Team (2021)) and constructed a network of those positive pairwise correlations above a threshold value of 0.3 to focus on the strongest correlations as suggested by Friedman and Alm (2012).

Lastly, we fit a community model to our metabarcoding data through spatiotemporal topic modeling (Girdhar et al., 2014). This method identifies and precisely describes clusters of taxa that are observed together regularly within our time series. We refer to these clusters as “co-occurrence communities”, which can be useful for decomposing our taxonomic focal groups (small phytoplankton and chlorophytes) into ecologically meaningful parts. Spatiotemporal topic modeling has previously been applied to time series of plankton image data (Kalmbach et al., 2017; San Soucie et al., 2024), but because it has not often been used to analyze genetic data, we briefly discuss the premise and benefits of the method below.

Spatiotemporal topic modeling is an extension of Latent Dirichlet Allocation that incorporates information about the distance in time and/or space between samples (Girdhar et al., 2014; Girdhar and Dudek, 2015). It is a tool designed to assess co-occurrence within high-dimensional, sparse, and compositional data. The underlying approach was developed primarily for text analysis, which, like metabarcoding data, involves large numbers of unique categorical observations that vary greatly in their relative abundance. That is, some words in text analysis and some ASVs in metabarcoding are common while others are rare. In their original use, topic models assign each word in a text with an underlying topic label (e.g., the subject of a document) that best explains the appearance of that word (Blei et al., 2003). A text can then be described by the mixture of subjects it includes. For our application, we used the same algorithm to describe each of our samples as a mixture of underlying “co-occurrence communities”. Reads were assumed to be sampled at random from the weighted mixture of cooccurrence communities present at each time point, where each co-occurrence community has its own latent distribution of taxa that are regularly observed together. The best-fit model consists of the distributions of taxa within each cooccurrence community as well as a time series of the distribution of co-occurrence communities within each sample. The results of spatiotemporal topic modeling are tied to complete descriptions of the community at each time point, and are thus more readily interpretable than clusters identified through PCA or within correlation networks.

In addition to being well suited to amplicon sequencing data in general, spatiotemporal topic modeling has four notable features that make it particularly appropriate for application to our time series: 1) Individual samples can include mixtures of co-occurrence communities that overlap in time, allowing us to consider gradual changes that might be resolved at our monthly sampling resolution.

2) Samples do not need to be regularly spaced in time, so we do not need to average observations or interpolate to fill gaps in our time series. 3) Information about the distance between samples (in time) is retained and used in the clustering process. A solution is considered more likely if it has fewer sudden changes in the prevalence of a co-occurrence community. Lastly, 4) a single taxon can contribute to multiple communities with equal or unequal probability. That is, taxa that are poorly resolved or that participate in multiple ecological associations can be present within multiple clusters. The last two benefits are not possible within the clustering that can be performed through PCA.

For this work, we used species-level assignments to fit models with two cooccurrence communities in order to compare to our PCA results. As we did for PCA, we conducted this analysis separately on the small phytoplankton and Chlorophyta focal groups. For each, the model assumes that both the distribution of co-occurrence communities within each sample and the distribution of taxa within each co-occurrence community are Dirichlet distributions. We used hyperparameter values of 0.1 and 0.5 for these two distributions, respectively. Previous work applying topic models to plankton observational data—and our own exploration of hyperparameter space—found community assignments to be qualitatively insensitive to values between 0 and 1 (San Soucie et al., 2024). We compared the topic model outputs to the pairwise relationships identified in SparCC and to the outputs of the PCA.

## 3 Results

### 3.1 Taxonomic composition of small phytoplankton

The small phytoplankton group is composed primarily of members of the phyla Chlorophyta (49% of reads), Heterokontophyta (23%), and Haptophyta (14%). Of these, Chlorophyta contribute the largest proportion of the reads measured at MVCO and account for 33% of the ASVs assigned at the division-level (Fig. S1B). The Chlorophyta ASVs are dominated by Mamiellophyceae, followed by Pyramimonadophyceae and Trebouxiophyceae (Fig. S1D). The Heterokontophyta are slightly more diverse than Chlorophyta in terms of ASV richness (856 vs 742 ASVs) but accounted for a smaller proportion of the reads. Within the Heterokontophyta, Bacillariophyta and Pelagophyceae contribute the largest fractions of reads. At the genus level, *Micromonas* is the most abundant genus across all samples, followed by *Bathycoccus* and *Picochlorum* (Fig. 1). Within individual samples, *Micromonas* and *Picochlorum* are most often the dominant taxa present (Fig. 1C). The Shannon diversity index calculated from the genus-level taxonomic assignments has an interquartile range of 2.12-2.87 for the small phytoplankton and 1.32-1.92 for the chlorophytes. These values are relatively consistent across the year but are lowest when *Picochlorum* or *Phaeocystis* make up the largest number of reads within a sample (Fig. S2).

**Fig. 1:**
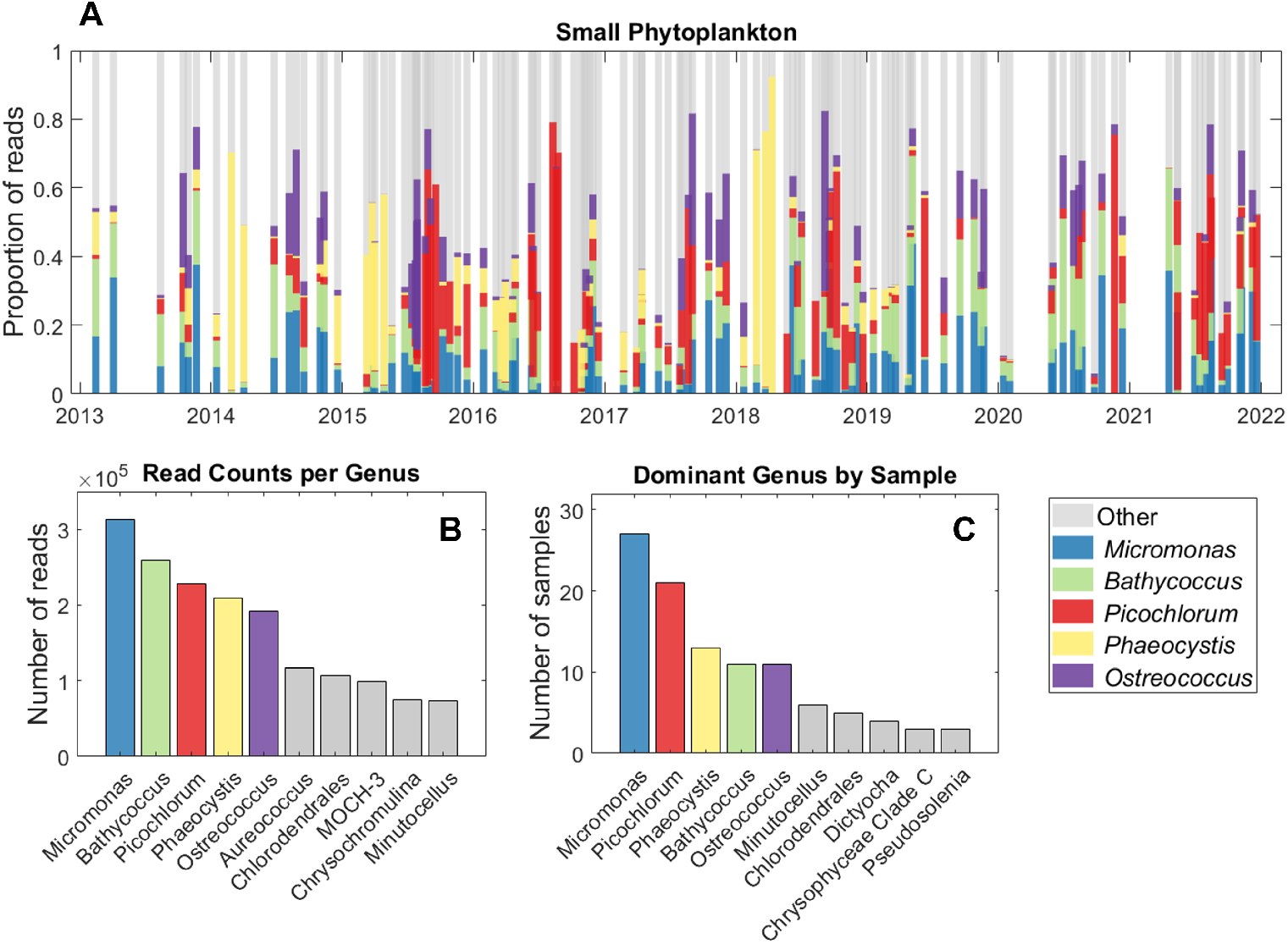
A) Timeline of the taxonomic composition of small phytoplankton at MVCO divided by genus. Highlighted are the 5 genera with the greatest contributions to B) the total number of reads in the time series. These genera are also dominant in terms of C) the number of samples in which each genus makes up the largest proportion of reads. Only the first 10 genera are included in panels B and C.

### 3.2 Annual patterns in community composition

Roughly 30% of the genetic variability in our dataset is explained by the first two components in our PCA (Fig. 2). While there are no distinct clusters within the PCA for either the small phytoplankton or Chlorophyta focal groups, there are gradual and repeated progressions through the coordinate space (Fig. 2). In both the small phytoplankton and chlorophyte analyses, principal component 1 (PC1) follows a seasonal pattern, with higher values in summer and lower in winter.

**Fig. 2:**
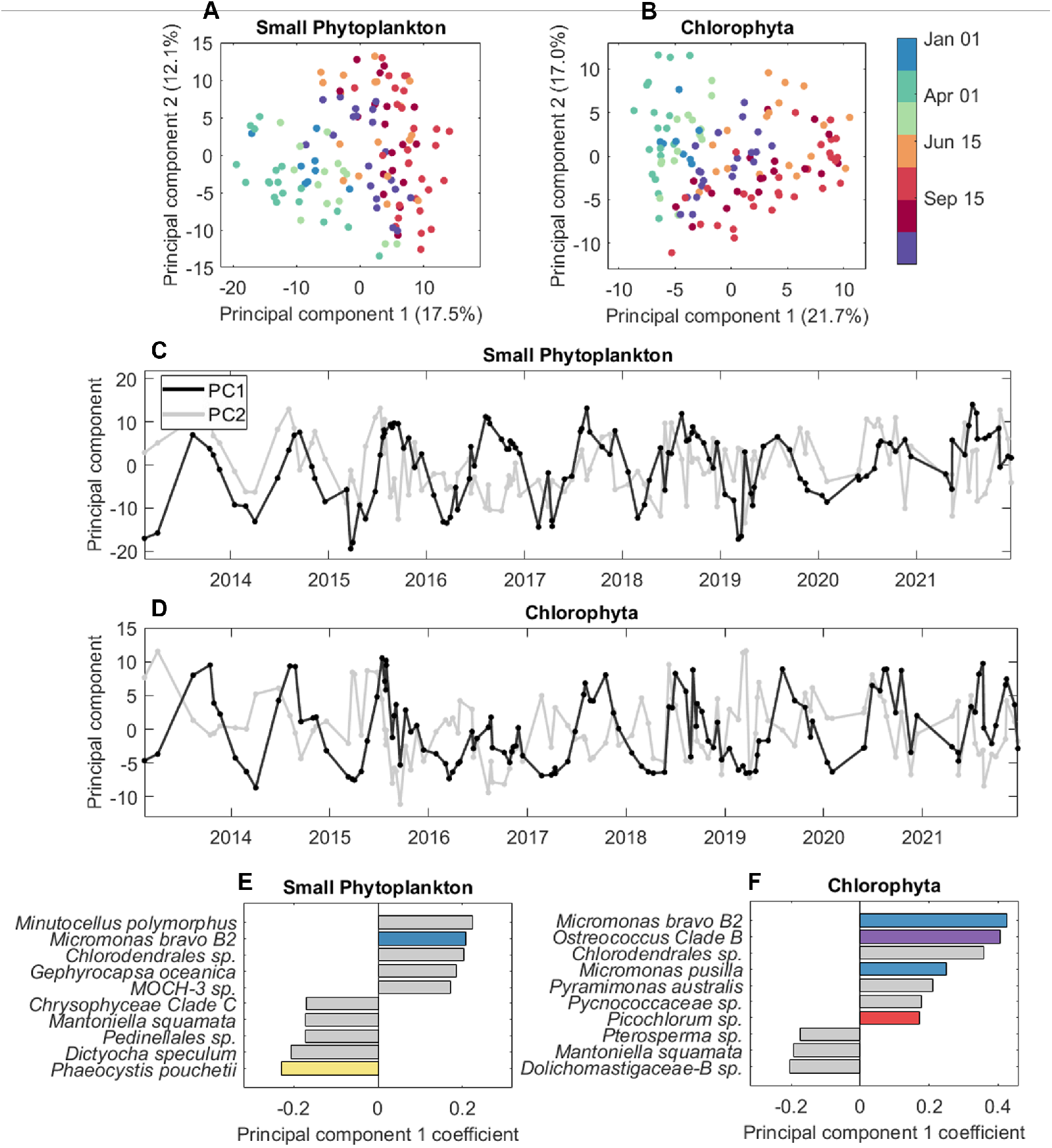
Principal component analysis of species-level assignments within two subsets of the metabarcoding data: the small phytoplankton and the chlorophytes. A.B) Each sample is plotted in two dimensional space according to the first two principal components. The percent of variance explained by each component is indicated in parentheses in the axes labels. Points are colored according to the time of year at which the samples were collected from MVCO. C,D) The first two principal component values in (A,B) plotted directly against sample date as a time series. E,F) The 10 species-level taxonomic assignments with greatest magnitude contributions to principal component 1 for each PCA. Taxa within the most abundant genera identified in Fig. 1 are colored accordingly.

Many environmental variables also have annual periodicity at MVCO, including temperature, salinity, and nutrient availability (Fig. S3). Seawater temperature is significantly correlated with PC1 values for both Chlorophyta (Pearson’s *R* = 0.75, *p <* 0.001) and small phytoplankton (Pearson’s *R* = 0.82, *p <* 0.001; Fig. S4). Phosphate concentration is also correlated with PC1 to a lesser extent (Fig. S4). When comparing cubic regressions trained on subsets of the data, temperature and day-of-year are equally good predictors of the values of PC1 for each group of phytoplankton: neither model significantly outperforms the other.

Finally, comparing both PCA results to our flow cytometry time series, we find that variability in the Chlorophyta more closely aligns with the abundance of phytoeukaryote cells. Over the annual cycle, PC1 for the small phytoplankton follows a smooth curve resembling the annual cycle in temperature (Fig. 3 A-B). In contrast, PC1 for Chlorophyta increases rapidly in the spring and decreases gradually in the fall, more similar to the pattern in phytoeukaryote concentration measured by flow cytometry (Fig. 3 C-D). Despite the comparable shape, the two cycles are slightly offset, with the peak in Chlorophyta PC1 occurring around 40 days after the peak in cell abundance (Fig. 3). Across the 76 sampling dates where genetic and flow cytometry measurements are both available, Chlorophyta PC1 is significantly correlated with daily average phytoeukaryote concentration (*R* = 0.36, *p* = 0.002), whereas the correlation between cell concentration and PC1 for the small phytoplankton is not significant (*R* = 0.18, *p* = 0.12).

**Fig. 3:**
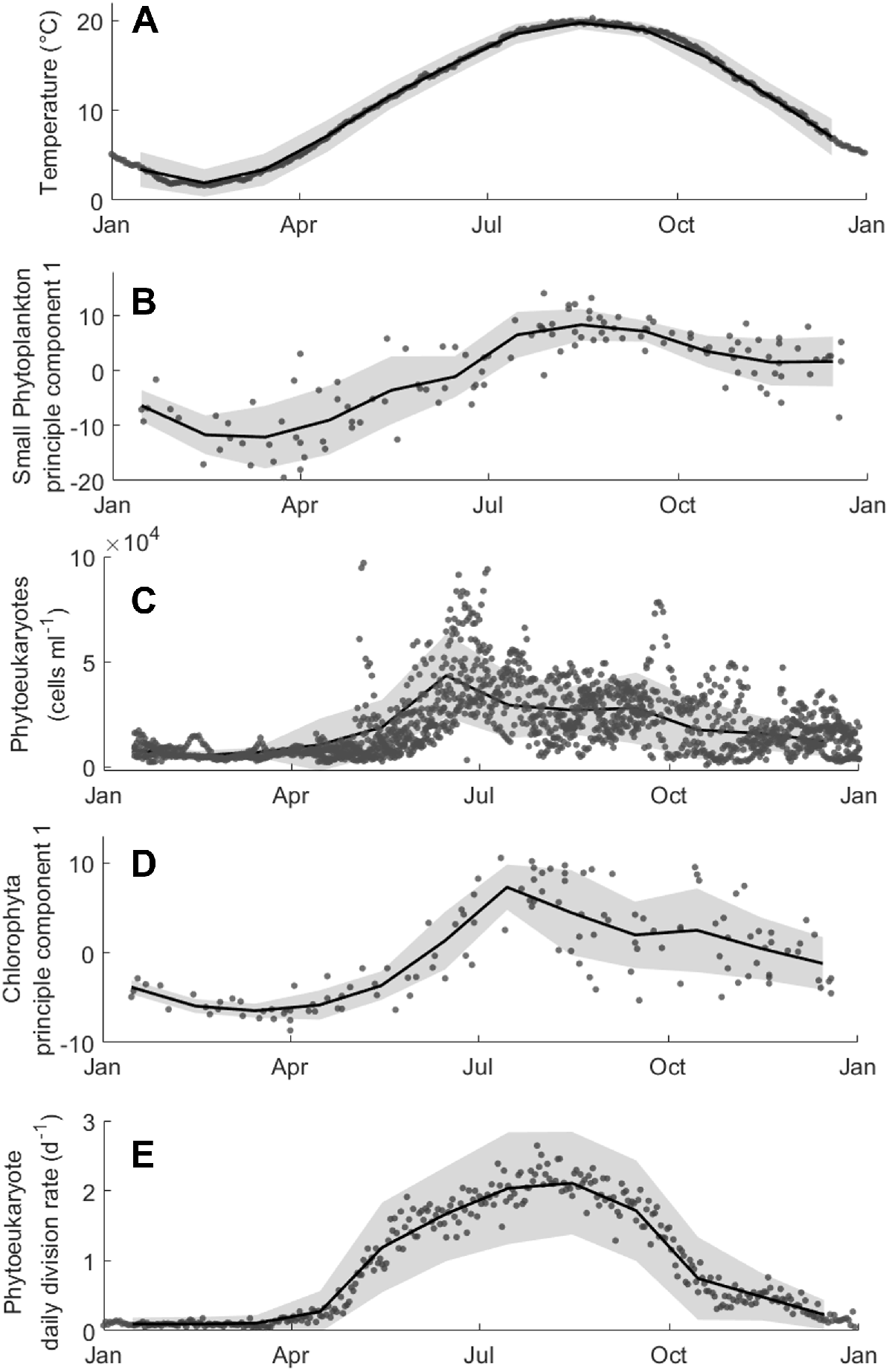
Annual cycles at MVCO. A) Seawater temperature at MVCO is well correlated with B) values of principal component 1 for the small phytoplankton group. C) The concentration of small phytoeukaryotes measured by FlowCytobot more nearly resembles the seasonal pattern in D) the principal component for Chlorophyta. E) Daily division rate estimates for the small eukaryotic phytoplankton assemblage reprinted from Fowler et al. (2020). In each panel, gray points indicate daily values, with the black line and shaded region showing the mean and standard deviation for each month of the year averaged across the time series (2013-2021 for A-D; 2003-2019 for E).

### 3.3 Co-occurring groups cluster by time of year

When clustering data into two co-occurrence communities, the best-fit model consists of one community that is observed most in summer and one observed most in winter (Fig. 4). At the species level, *Bathycococcus prasinos* and *Micromonas commoda* A2 are the most likely to be sampled from the winter community and *Picochlorum* sp. is the most likely to be sampled in the summer. Note that while the algorithm that identifies this model does incorporate the distance in time between samples, it does not have any information about environmental variables, nor would it have any inherent bias for an annual periodicity. The time-series outputs of the spatiotemporal topic modeling for chlorophytes and small phytoplankton differ only slightly (Fig. 4A, S5A), with the inferred cooccurrence communities in both analyses largely made up of Chlorophyta (Fig. 4B,C).

**Fig. 4:**
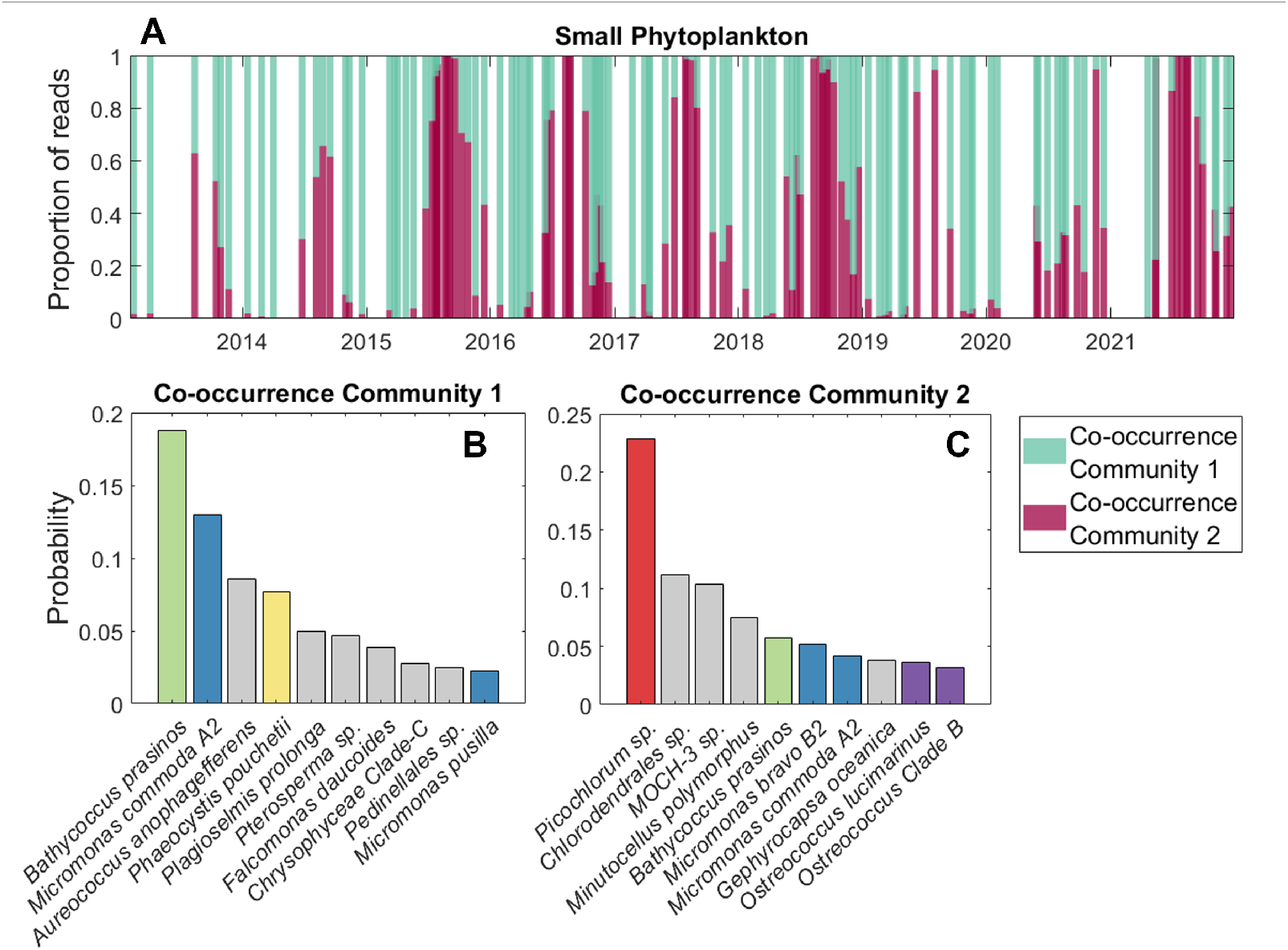
Timeline of the taxonomic composition of small phytoplankton at MVCO summarized by the best-fit topic model with two co-occurrence communities. A) Proportion of reads in each sample attributed to each of the two co-occurrence communities. B-C) The 10 taxa with the greatest probability of being sampled from co-occurrence communities 1 and 2, respectively. Taxa within the dominant genera identified in Fig. 1 are colored accordingly. Note the improved interpretability of the time series compared to Fig. 1.

SparCC analysis identifies 22 chlorophyte taxa with strong positive correlations to one another. These relationships cannot be separated into distinct networks, such that every node is connected indirectly to every other (Fig. 5). The correlations identified reinforce the associations between small phytoplankton taxa evident in the PCA and co-occurrence communities. For example *Micromonas bravo* B2 is correlated with *Chlorodendrales, Picochlorum*, and *Ostreococcus* clade B, which have all been identified by the other methods as more prevalent in summer. *Bathycoccus prasinos* and *Micromonas commoda* A2 are positively correlated, as would be expected by their prevalence within the inferred winter co-occurrence community, and *Pterosperma, Mantoniella squamata*, and *Dolichomastigaceae* B, the three negative contributors to PC1 for Chlorophyta, are all correlated as well.

**Fig. 5:**
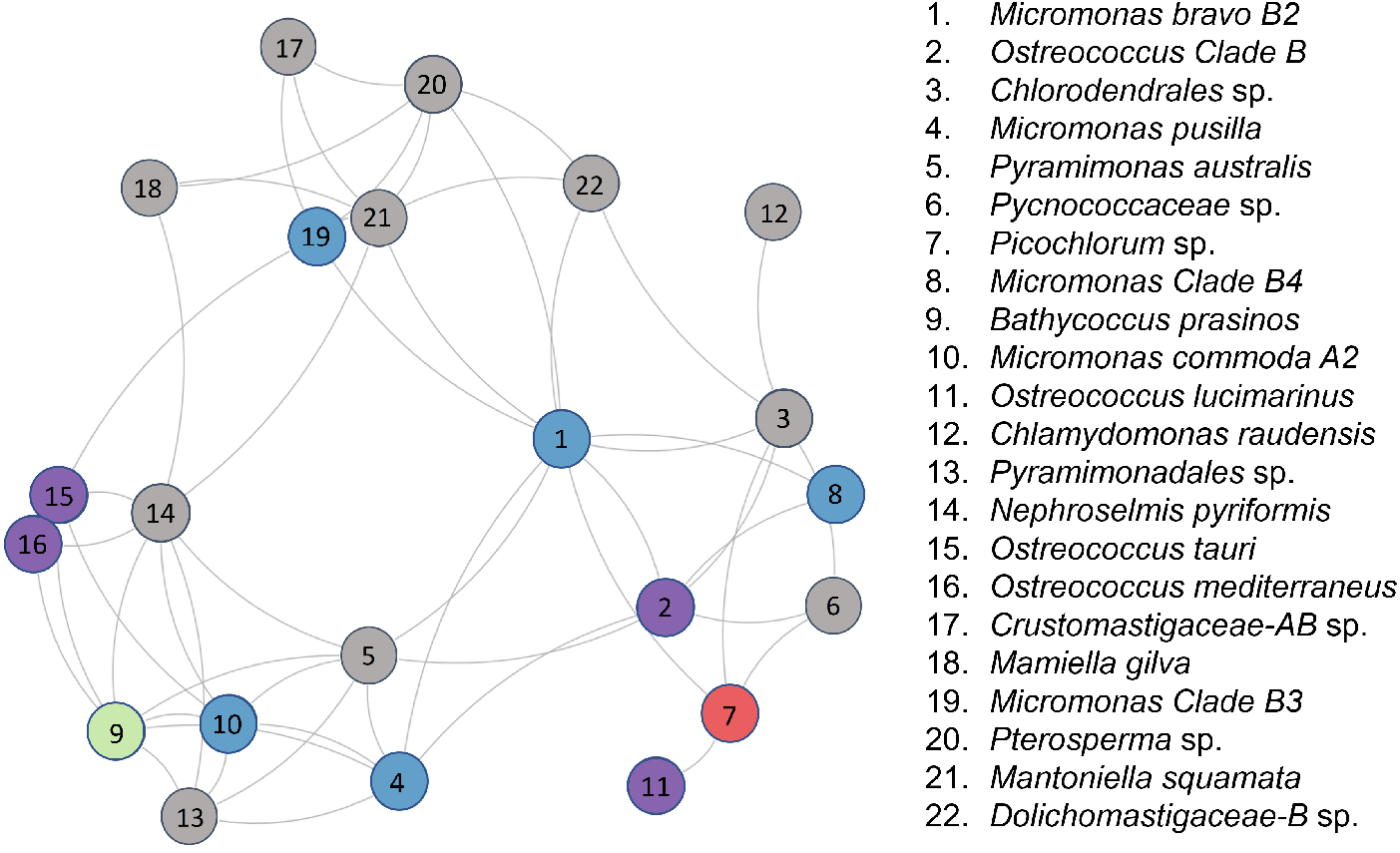
Network of associated chlorophyte taxa, as identified by SparCC analysis. Positive pairwise correlations greater than 0.3 are drawn as gray lines connecting species-level taxonomic assignments. Taxa that were not identified as having significant correlations with any others are not drawn in the network. Nodes are numbered according to the relative contribution of each taxon to principal component 1, shown in Fig. 2, and colored by genus, as in Fig. 1.

## 4 Discussion

### 4.1 Key contributors to the small phytoplankton assemblage

We have presented an analysis of genetic samples from 125 days over a 9-year period in order to characterize the taxonomic composition of the pico- and nanosized eukaryotic phytoplankton in the well-studied Northeast U.S. Shelf ecosystem. Within our data set, we find that Chlorophyta make up the majority of the reads identified as small phytoplankton. *Micromonas* and *Bathycoccus* are the dominant genera by total read count and are detectable at the site year-round. *Ostreococcus* is also prevalent at MVCO in all seasons at slightly lower read counts. Species of *Picochlorum* and *Phaeocystis* appear in the time series more episodically, making up large portions of the reads in summer and winter samples, respectively.

The prevalence of Mamiellophyceae at our site is consistent with many coastal locations around the world (Tragin and Vaulot, 2018), including but not limited to the Northwest Pacific (Lin et al., 2017), South East Pacific (Rii et al., 2016), English Channel (Not et al., 2004), Mediterranean (Zhu et al., 2006), and Indian Ocean (Not et al., 2008). The particular dominance of *Micromonas* and *Bathycoccus* is also widespread, prompting some authors to look specifically for evidence of niche partitioning between these two genera (Joli et al., 2017). One complicating factor when looking for such patterns is the potential for unresolved diversity within the genera. For example, Limardo et al. (2017) found that multiple ecotypes exist within the species *Bathycococcus prasinos*, each with distinct temperature and nutrient preferences. In our analysis, *Bathyoccocus* sequences can only be assigned to a single group. However, multiple species of *Micromonas* can be distinguished. We find *Bathycoccus* reads correlate with reads assigned to *Micromonas commoda* A2 and *Micromonas pusilla*, but not with other *Micromonas* groups. While these exact species assignments are not certain, the differences suggest niche differentiation at the species level.

*Ostreococcus* is a third genus of the class Mamiellophyceae that is frequently observed in marine samples. Off the coast of southern California, for examples, quantitative PCR revealed that *Ostreococcus* was present in almost every sample and made up as much as 70% of the picophytoeukaryotes (Countway and Caron, 2006). At MVCO, *Ostreococcus* was present in 107 out of our 125 samples. The somewhat lower total read count compared to *Bathycoccus* and *Micromonas* may reflect lower abundance, or may be due to its lower gene copy number (Zhu et al., 2006) and small genome (Palenik et al., 2007). It is also likely that *Ostreococcus* reaches higher densities further offshore or below the surface waters we sampled, as members of the genus have been seen to be particularly dominant at the base of the mixed layer and along upwelling coasts where nutrients are more readily available (Countway and Caron, 2006; Demir-Hilton et al., 2011; Collado-Fabbri et al., 2011; Rii et al., 2016).

While present throughout the year at MVCO, *Picochlorum* tends to dominate samples in the summer (Fig. 1). Globally, *Picochlorum* has been seen to be the most abundant trebouxiophyte in temperate waters (Tragin and Vaulot, 2018). Species in this genus have been found to be halo-tolerant and to divide rapidly at temperatures above 25^*°*^ C, which is well above the typical summer temperatures at MVCO (Henley et al., 2004; Stawiarski et al., 2016). These traits may render *Picochlorum* more likely than other small phytoplankton taxa to bloom at MVCO under specific summer conditions. Additionally, certain cultured *Picochlorum* species are documented mixotrophs (Pang et al., 2022), which is important to consider when studying the planktonic food web. A high degree of mixotrophy in the small phytoplankton, for example, might explain the decoupling between growth and grazing measured in summer incubation experiments on the Northeast U.S. Shelf (Marrec et al., 2021).

*Phaeocystis* is a second genus that sporadically dominates the samples in our time series, but, unlike *Picochlorum*, is more often observed in winter. *Phaeocystis* is a colonial organism that undergoes intense blooms in many parts of the world (Veldhuis et al., 1986; Smith et al., 2021). Notably, individual *Phaeocystis* cells are on the larger end of cell volumes included in our flow cytometry measurements (4-10 *µ*m) and the colonies are too large to be included (up to 2-8 mm) (Throndsen, 1997). Thus, despite its large genetic signal, *Phaeocystis* may not always be measured by FlowCytobot when present. Our flow cytometry may, however, be capturing free-living *Phaeocystis* cells or those shed by colonies during bloom senescence. This might produce the bimodality in small eukaryote cell volume that is observed repeatedly in spring at MVCO (Fowler et al., 2020) (Fig. S6). Direct microscopy or cell sorting would be needed for confirm whether these larger cells are in fact *Phaeocystis*. Prompted by a particularly dramatic *Phaeocystis* bloom in spring of 2018, Smith et al. (2021) compared genetic data from MVCO to a detailed reference collection of *Phaeocystis* sequences. They found the best match to be *Phaeocystis pouchetii*. That information is reflected in the species labels of this paper (e.g., Fig. 2), but we note that the automatic assignment from our DADA2 pipeline is *Phaeocystis antarctica*. Such an error highlights the limits of this analysis: assignments may be imprecise at the species level. That said, variability observed within the genetic data will reflect true variability in the reads regardless of their classification.

### 4.2 Temporal variability in community composition

The composition of the small phytoplankton community at MVCO undergoes smooth seasonal transitions that are repeated in every year of our time series. For both the small phytoplankton and Chlorophyta groups in our analysis, the principal component that explains the most variability in the data follows an annual cycle (Fig. 2B), and the best-fit topic model identifies communities that group into seasons. The largest differences in the community are observed roughly be-tween February and August, with gradual transitions between these extremes during the spring and fall. Our monthly sampling resolution is sufficient to capture this pattern of succession over the annual cycle.

Because multiple environmental variables change seasonally on the Northeast U.S. Shelf, it is hard to identify a specific driver of the observed changes in community composition. In other regions of the world, picoeukaryote communities change in response to annual trends in river runoff (Jing et al., 2010), wind-driven upwelling (Collado-Fabbri et al., 2011), and temperature (Li et al., 2017). We find that seawater temperature is significantly correlated with PC1 for both the small phytoplankton and Chlorophyta, but that temperature is no stronger a predictor of the value of PC1 than day-of-year. While release from temperature limitation is a driver of the spring bloom of both *Synechococcus* and picoeukaryotes at MVCO (Hunter-Cevera et al., 2016; Fowler et al., 2020), it remains possible that some small phytoplankton groups are responding to their own endogenous rhythms (Lambert et al., 2018), or indirectly to the environment via the activity of other taxa.

Despite changes in the prevalence of individual taxa between samples, we find total diversity metrics within the small phytoplankton community to be relatively stable (Fig. S2). This suggests a degree of robustness at the aggregate level, even if environmental change were to shift conditions in favor of certain taxa over others. For example, if regional warming were to lead to shorter windows of favorable conditions for the winter community, a diversity of small phytoplankton would likely persist in the warmer conditions. Biomass-specific primary production has been seen to be consistent across distinct picoeukaryote communities (Grob et al., 2011). However, more detailed knowledge is needed to understand the ecological consequences of losing or replacing any individual small phytoplankton taxa.

Correlations identified through SparCC can help guide more detailed ecological study by pointing to important pairwise relationships within the plankton. Strong correlations could indicate overlapping niches and heightened competition or facilitation, but need to be confirmed through targeted effort. The correlation network of Chlorophyta (Fig. 5) shows taxa that are correlated over time. Within the network, we can see a signature of the gradual transitions in community composition that are evident in our other analyses. Because the seasonal succession at MVCO is gradual, sequentially blooming taxa create paths in the network between taxa that do not overlap in their peak occurrence. Even taxa which are present at opposite times of year (e.g., *Micromonas bravo* B2 and *Bathycoccus prasinos*) are connected in the network via mutual positive correlations with intermediate taxa. Through further interrogation, we may be able to discern paths by which winter microbial dynamics affect summer dynamics and vice versa.

### 4.3 Comparing taxonomic composition to flow cytometry measurements

During the annual spring bloom at MVCO, small phytoeukaryote concentration increases from *∼*300 cells ml^*−*1^ to over 50,000 cells ml^*−*1^ (Fig. 3; Fowler et al. (2020)). The sustained diversity of the small phytoplankton community over the annual cycle suggests that the spring bloom is not monospecific, but either a shared response of a group or the result of relatively even turnover between the dominant taxa at different times of year. Comparing cell abundance to the variability in community composition, we find that cell abundance is significantly correlated with the Chlorophyta, as indicated by the first principal component of our PCA (Fig. S4). While there is no way to definitively show that the sequenced taxa are those cells we are measuring with flow cytometry, this result is a strong indication that the chlorophyte taxa contributing to PC1 are part of the spring bloom of small phytoplankton at MVCO.

The annual cycle in PC1 for chlorophytes very nearly reflects the annual pattern in cell density measured by flow cytometry. Both metrics increase dramatically in the spring and decrease gradually over the fall (Fig. 3; Fig. S7). Interestingly, the annual peak in PC1 occurs after the peak in cell abundance. Community turnover after the bloom’s peak suggests a mechanism for a surprising finding from earlier work: that picoeukaryote division rate reaches its maximum roughly 60 days after the average peak in cell concentration (Fig. 3E; Fowler et al. (2020)). Those division rates are representative of the entire assemblage, meaning that the presence of slower growing cells would bring down the community average. Since PC1 effectively indicates the distance between the Chlorophyta community and the typical community in winter, the time lag we report might be the result of winter taxa that remain in the community until after the spring bloom reaches its peak. As slower growing strains are removed from the community, the measured division rate would increase, even if all strains were growing at their individual maxima throughout the bloom. If this is the case, the increase in division rate is an emergent result of top-down control on the small phytoplankton. This intriguing possibility underscores the importance of measuring changes in community composition to provide context for aggregate dynamics. By combining metabarcoding data with high-resolution quantitative tools, we can gain a more complete picture of the ecology within diverse microbial communities.

## Acknowledgements

The authors would like to thank Taylor Crockford, Emily Brownlee and Kristen Hunter-Cevera for collecting many years of genetic samples from MVCO, as well as Rob Olson and Alexi Shalapyonok for generating and maintaining the flow cytometry time series. Thank you to Ann Tarrant for allowing the lead author to use her lab space for DNA extraction, to Tim Shank for use of his microvolume spectrophotometer, to Neel Aluru for the well plate, and to Cory Berger for teaching the lead author the extraction protocol. This research was made possible by National Science Foundation (#OCE-1655686, #OCE-2322676, and #OCE-1434440), the Simons Foundation (561126, 1188720, and LS-FMME-00008380), and the Grassle Student Fellowship Fund.

## Author Contributions

BS, RG, MN, and HS conceptualized research and acquired funding. BS, RG, EP, and HS contributed to data curation and investigation. BS, RG, EP, YG, and HS developed methodology. RG and HS performed data validation. BS, EP, YG, and HS developed software. BS and HS provided resources. BS, MN, and HS contributed to project administration and supervision. BS drafted initial manuscript and created visualizations. All authors reviewed and edited the final manuscript.

## Data Availability Statement

Sequence data have been deposited with links to BioProject accession numbers PRJNA504617 and PRJNA1170325 (https://www.ncbi.nlm.nih.gov/bioproject/PRJNA504617, https://www.ncbi.nlm.nih.gov/bioproject/1170325).

Flow cytometry data have been deposited to the Environmental Data Initiative (https://doi.org/10.6073/pasta/99f139f2dbdea0b09dba02dfead1ffdc).

