## Supplementary Figures S1-S7 for "Small phytoplankton community composition cycles annually with a coastal bloom"

### Supplementary information for " Small phytoplankton community composition cycles annually with a coastal bloom"

[4] Applied Ocean Physics and Engineering
Woods Hole Oceanographic Institution
Woods Hole, MA 02543

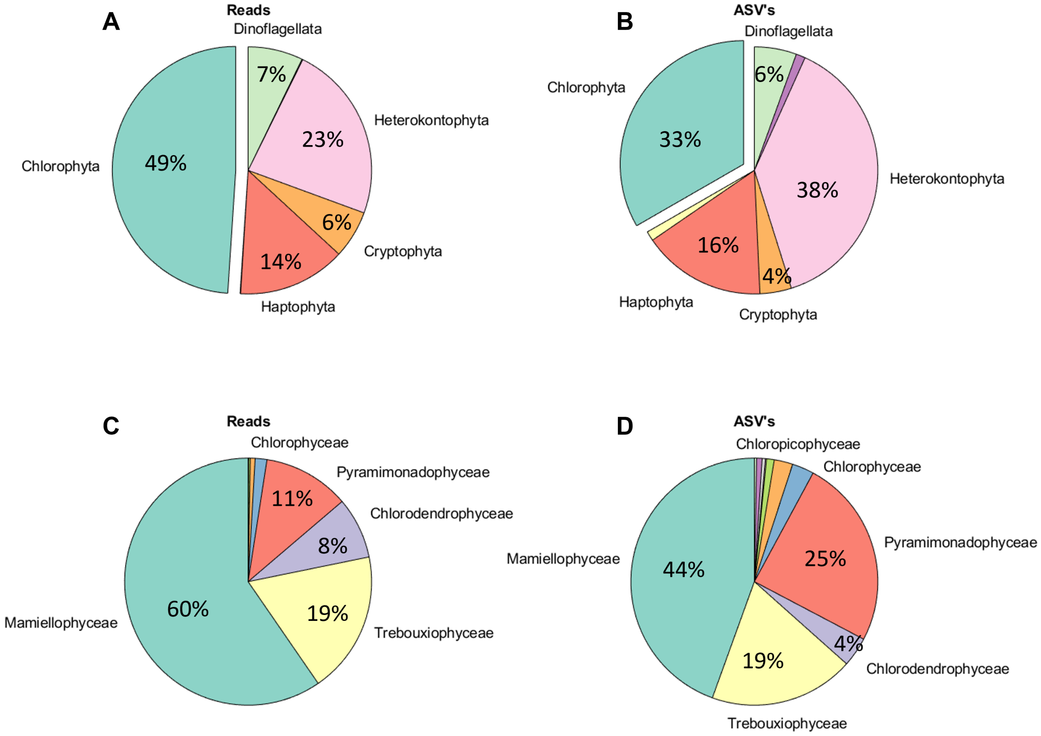


Fig. S1: Contribution to read counts (left) and diversity (right) of A-B) phyla within the small phytoplankton focal group, and C-D) classes within the Chlorophyta component of the small phytoplankton group. Note that the Chlorophyta analyzed are a strict subset of the small phytoplankton and do not include macroalgae, Ulvophyceae. Unlabelled wedges make up 2% or less of the total.

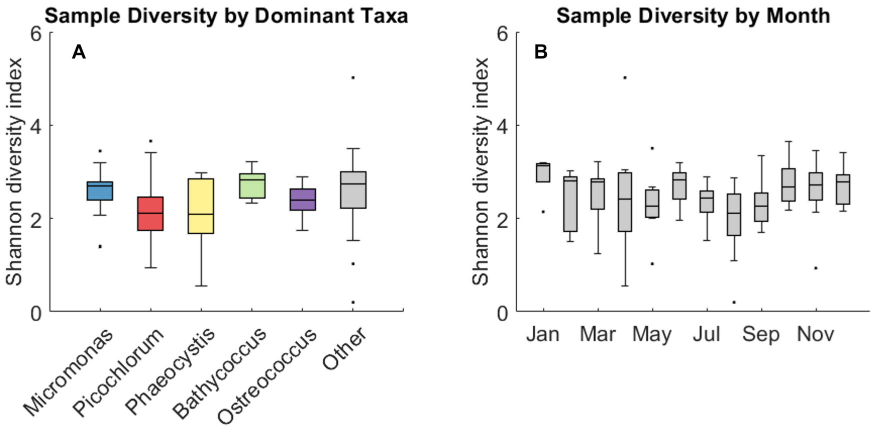


Fig. S2: Shannon diversity indices of small phytoplankton in metabarcoding samples grouped according to A) the genus with the largest proportion of reads in a sample and B) the month sampled. Horizontal lines indicate median, boxes indicate interquartile range, and dots indicate values identified as outliers based on interquartile method.

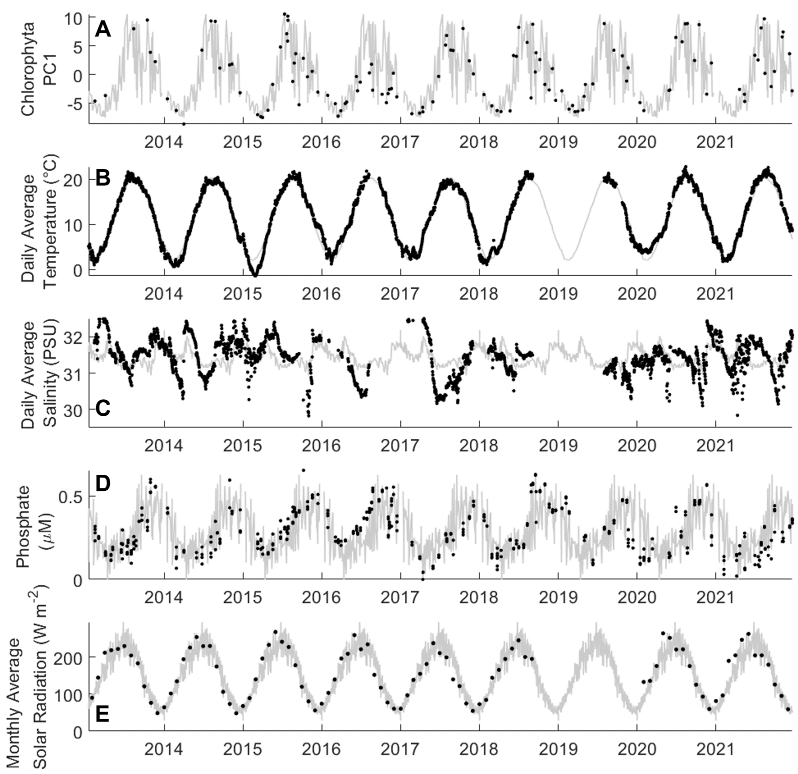


Fig. S3: Time series of seasonality at MVCO. A) Value of principal component 1 for Chlorophyta as shown in Fig. 2. Daily average seawater B) temperature and C) salinity measured continuously at MVCO. D) Discrete measures of dissolved phosphate from the upper 10 meters of the water column. E) Monthly averaged incident solar radiation. In all panels, gray line is the climatology, or average annual cycle over the time series at daily resolution.

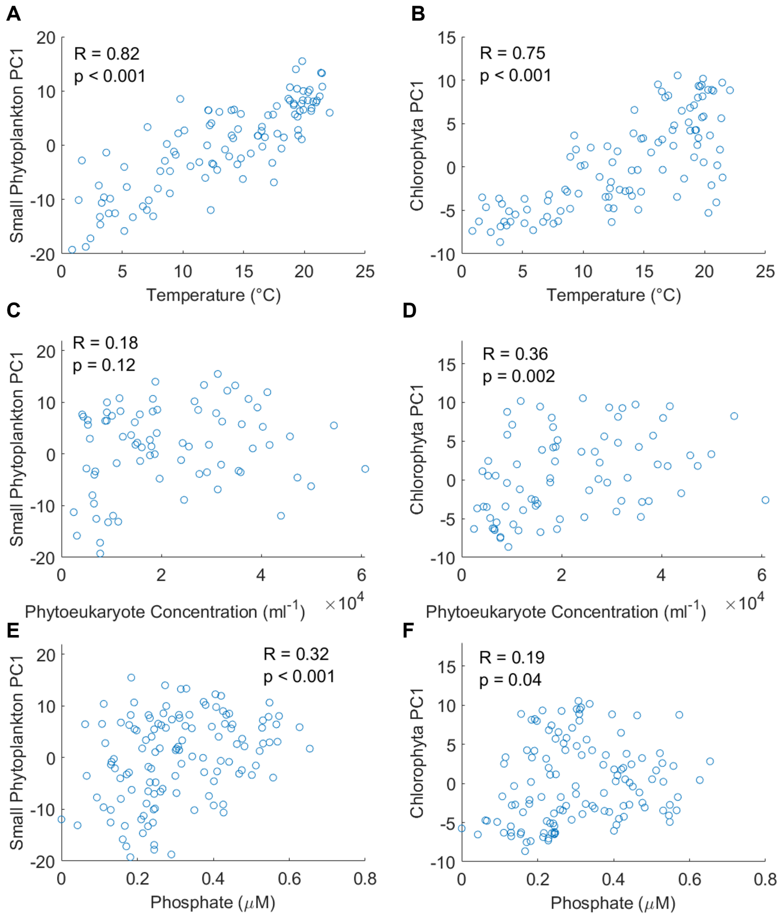


Fig. S4: Relationship between small phytoplankton compositional variability and A-B) seawater temperature, C-D) phtyoeukaryote abundance measured by Flow-Cytobot, and E-F) dissolved phosphate concentration. Y-axes show values of PC1 for the small phytoplankton focal group (left) or Chlorophyta focal group (right). Pearson correlation coefficients (R) and associated p-values are indicated within each panel.

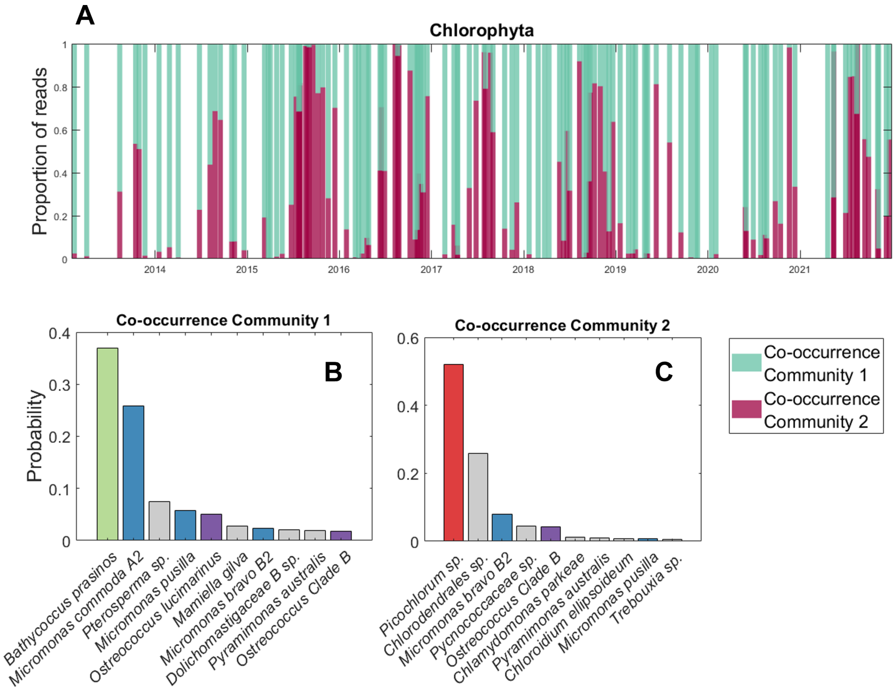


Fig. S5: Timeline of the taxonomic composition of Chlorophyta at MVCO summarized by the best-fit topic model with two co-occurrence communities. A) Proportion of reads in each sample attributed to each of the two co-occurrence communities. B-C) The 10 taxa with the greatest probability of being sampled from co-occurrence communities 1 and 2, respectively. Taxa within the dominant genera identified in Fig. 1 are colored accordingly. Note that this model was fit to the Chlorophyta taxa independent of the model fit to small phytoplankton presented in Fig. 4.

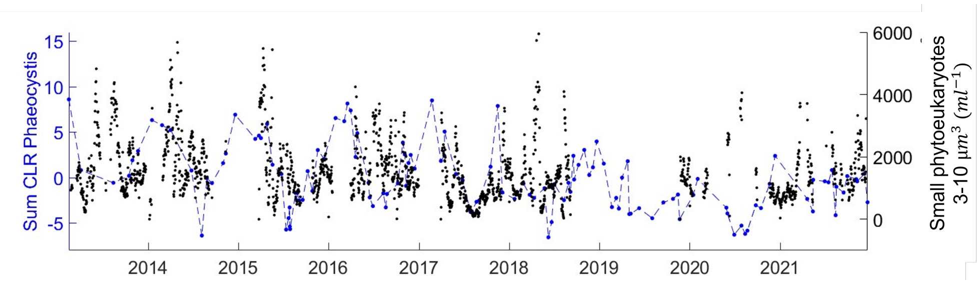


Fig. S6: Centered log ratio (CLR) of all reads assigned to the genus *Phaeocystis* (blue circles and dotted line) overlain on time series of the daily average concentration of cells between 3 and 10 $\mu$m in diameter measured by FlowCytobot (black circles). *Phaeocystis* CLR peaks regularly in winter and in multiple years nano-eukaryote concentration increases sharply as *Phaeocystis* CLR begins to decline.

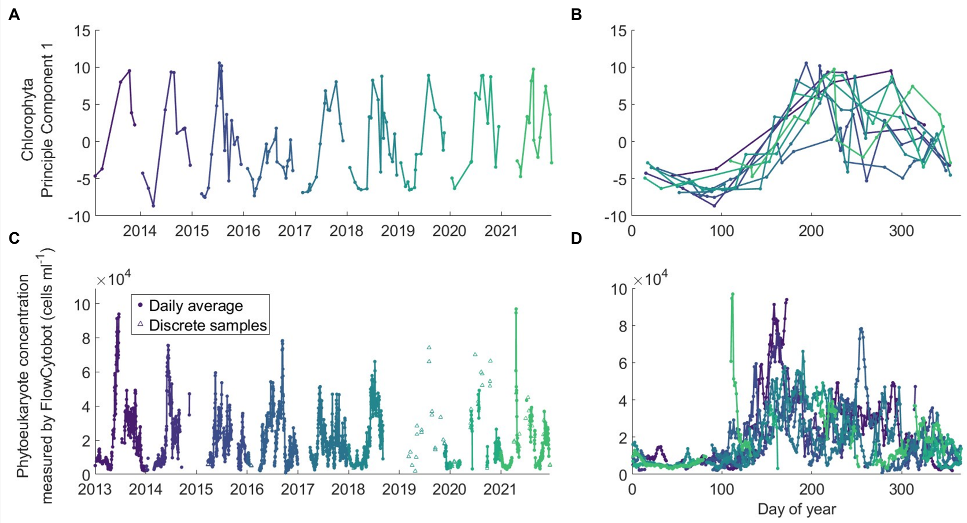


Fig. S7: Correspondence between genetic and flow cytometric data. Values of principal component 1 from the Chlorophyta focal group plotted against A) date and B) day of year. Daily average in situ phytoeukaryote concentration (circles) and individual phytoeukaryote concentration measurements from discrete samples (triangles) plotted against C) date and D) day of year. Points and lines in all panels are colored by year sampled.
